# Feasibility of Topological Data Analysis for event-related fMRI

**DOI:** 10.1101/457747

**Authors:** Cameron T. Ellis, Michael Lesnick, Gregory Henselman-Petrusek, Bryn Keller, Jonathan D. Cohen

**Affiliations:** Department of Psychology, Yale University, New Haven, CT, USA; Princeton Neuroscience Institute, Princeton University, Princeton, NJ, USA; Department of Mathematics, University at Albany, Albany, NY, USA; Intel Labs, Hillsboro, OR, USA

**Keywords:** topological data analysis, persistent homology, fMRI, simulation, event-related design, representation

## Abstract

Recent fMRI research shows that perceptual and cognitive representations are instantiated in high-dimensional multi-voxel patterns in the brain. However, the methods for detecting these representations are limited. Topological Data Analysis (TDA) is a new approach, based on the mathematical field of topology, that can detect unique types of geometric features in patterns of data. Several recent studies have successfully applied TDA to study various forms of neural data; however, to our knowledge, TDA has not been successfully applied to data from event-related fMRI designs. Event-related fMRI is very common but limited in terms of the number of events that can be run within a practical time frame and the effect size that can be expected. Here, we investigate whether persistent homology — a popular TDA tool that identifies topological features in data and quantifies their robustness — can identify known signals given these constraints. We use fmrisim, a Python-based simulator of realistic fMRI data, to assess the plausibility of recovering a simple topological representation under a variety of conditions. Our results suggest that persistent homology can be used under certain circumstances to recover topological structure embedded in realistic fMRI data simulations.

## Introduction

A fundamental construct in cognitive psychology and neuroscience is that of a representational space, within which knowledge is stored. A representational space is a set of dimensions along which items of a particular type are described, in which each dimension reflects features of items of that type (e.g., perceptual, semantic, evaluative, etc.). Thus, a central goal of both cognitive psychology and neuroscience is to characterize the dimensions that define representational spaces for different types of information (e.g., objects in the world, goals, classes of actions, etc.). Progress towards this goal obviously relies on methods that can identify the structure of such spaces. Cognitive neuroscience seeks evidence for such structure in patterns of neural activity.

Topological Data Analysis (TDA) holds promise for this effort, as an unbiased (that is, “exploratory”) method for identifying the latent structure in high-dimensional data. TDA uses ideas from the mathematical field of algebraic topology to study the shape (i.e., coarse-scale, global, non-linear geometric features) of data. For example, TDA seeks to detect clusters, holes, hollow voids, and tendrils in data (Carlsson, 2009). TDA has recently been applied productively to data from direct neuronal recordings. For instance, realistic models (Dabaghian, Mémoli, Frank, & Carlsson, 2012) and real data (Giusti, Pastalkova, Curto, & Itskov, 2015) suggest that the co-firing of hippocampal place cells contains topological structure. Other studies have shown that TDA can find structure in fMRI data (Bassett & Sporns, 2017). Examples of TDA being applied to fMRI data show that when a signal is periodic, TDA can capture it (Knyazeva, Orlov, Ushakov, Makarenko, & Velichkovsky, 2016), and TDA can describe the topological structure identified in the state transitions between tasks using other methods (Saggar et al., 2018). However, to our knowledge, TDA has not yet been demonstrated to directly identify structure in event-related fMRI data.

Event-related fMRI, in which discrete events are presented one after the other, is an important method for identifying mental representations. An example of a representation is the one you have of a landmark near your home (inspired by an experiment from Nielson, Smith, Sreekumar, Dennis & Sederberg, 2015). This representation has many dimensions (e.g., its location relative to your home, the type of place it is, how much you like it, how often you go there, etc.). Your representation of different landmarks around your home will have different coordinates in this representational space. To identify these representations, an event-related fMRI experiment could be carried out in which you are shown images of these landmarks one after the other, separated by a pause. Each image presentation is an event and will evoke a neural response. If these neural responses capture the spatial dimensions of these landmarks, then the physical distances in these landmarks will relate to the differences in neural representations. Neural responses might also capture perceptual or evaluative dimensions (e.g., their sensory features and/or how much you like each), in which case differences along these dimensions should relate to differences in neural representations. With this logic, it should be possible to identify patterns of activity in the brain that reflect differences between these landmarks along each of these dimensions; that is, their topological structure. This is a simple example but it applies to other types of representational spaces such as faces (Valentine, 1991), temporal relationships (Schapiro, Rogers, Cordova, Turk-Browne, & Botvinick, 2013), etc. Some fMRI studies have generated evidence for patterns of brain activity that reflect the metric properties inherent in a set of stimuli (Nau, Schröder, Bellmund, & Doeller, 2018; Schapiro et al., 2013), but these did not use TDA.

An appeal of TDA is that it has the potential to identify the topological structure of representational spaces (e.g., contiguity relationships) without requiring that affine properties (i.e., exact metric relationships) be preserved; for example, a rubber band preserves its topological identity even as it is stretched into various shapes. This may be important for identifying representational spaces in the brain, if these have topological structure that goes beyond strict metric relationships — something that standard methods would find harder to detect.

In this work, we focus on the application of persistent homology (Zomorodian & Carlsson, 2005), one of the most widely studied and applied TDA tools, to the analysis of representations in event-related fMRI. In order to evaluate the potential usefulness of persistent homology for identifying the structure of representational spaces in the brain, we simulated an event-related fMRI dataset by embedding a simple topologically-structured signal (a loop/ring) within a realistically noisy model of fMRI data. We tested different design parameters (e.g., number of event types, and samples per event type), as well as different levels of signal strength, to evaluate the extent to which persistent homology is able to recover the topological structure of the signal imposed on the noisy data.

## Methods

### Materials and Procedure

Our simulations were based on parameters extracted from raw fMRI data. We obtained the data from Schapiro and colleagues (2013). We chose to work with this data because many of its characteristics were representative of other fMRI studies (including its size: 20 participants, each with 5 runs); however, we note that some aspects of the design we simulated are distinct from the design of these authors (as described below). This data was collected on a 3T scanner (Siemens Allegra) with a 16-channel head coil and a T2* gradient-echo planar imaging sequence (TR=2s, TE=30ms, flip angle=90°, matrix=64×64, slices=34, resolution=3×3×3mm, gap=1mm). We used fmrisim (Ellis, Baldassano, Schapiro, Cai & Cohen, 2019) to simulate data with equivalent noise properties and embed a known topological signal into this realistic noise. The code is published at https://github.com/CameronTEllis/event_related_fmri_tda. An extensive description of fmrisim is outside of the scope of the current manuscript; we refer interested readers to Ellis and colleagues (2019) for a thorough description of how it works and analyses validating the accuracy of the simulations. Importantly, this package uses raw fMRI data to estimate noise parameters that can then be used to make a simulation with noise properties that approximate as closely as possible those in real data. This means that properties like the spatial structure, temporal variability and signal-to-noise ratio are similar between a simulation of a participant and the participant’s raw data. This can then be used to simulate data with a pre-specified (i.e., synthesized) signal embedded in it and observe “how much signal” is needed to observe statistically meaningful effects.

In this study, the noise parameters were estimated from the Schapiro and colleagues (2013) dataset and used to generate a noise volume. Figure S1 shows plots demonstrating the extent to which the simulated noise properties approximate real noise properties. These analyses show that the simulated participants have typical and appropriate noise properties. With these noise volumes, a pre-specified signal (a loop) was linearly added following the steps demonstrated in Figure S2. To explain these steps, the spatial structure of the signal was a set of N equi-spaced points on the circle, where N = 12, 15, or 18. We embedded these N points into high-dimensional voxel space via a fixed linear, Euclidean distance-preserving (orthonormal) transformation. Specifically, we embedded the N points into a region of interest (ROI) of 442 voxels (and only these voxels) spanning the left superior temporal gyrus, as well as the anterior temporal lobe and inferior frontal gyrus (based on the significant voxels in an analysis by Schapiro et al., 2013). Note that some of these voxels in the ROI were outside of the brain due to differences in brain anatomy between participants, so no signal was inserted into these voxels for that participant/run (mean: 381.0, range: 347-411). This process generated N vectors in a 442-dimensional subspace of voxel space, arranged in a circle. The embedding into the 442-dimensional space was chosen at random and the range of values were shifted to all be positive. Figure 1 shows schematically how the loop structure was embedded into the ROI.

**Figure 1.**
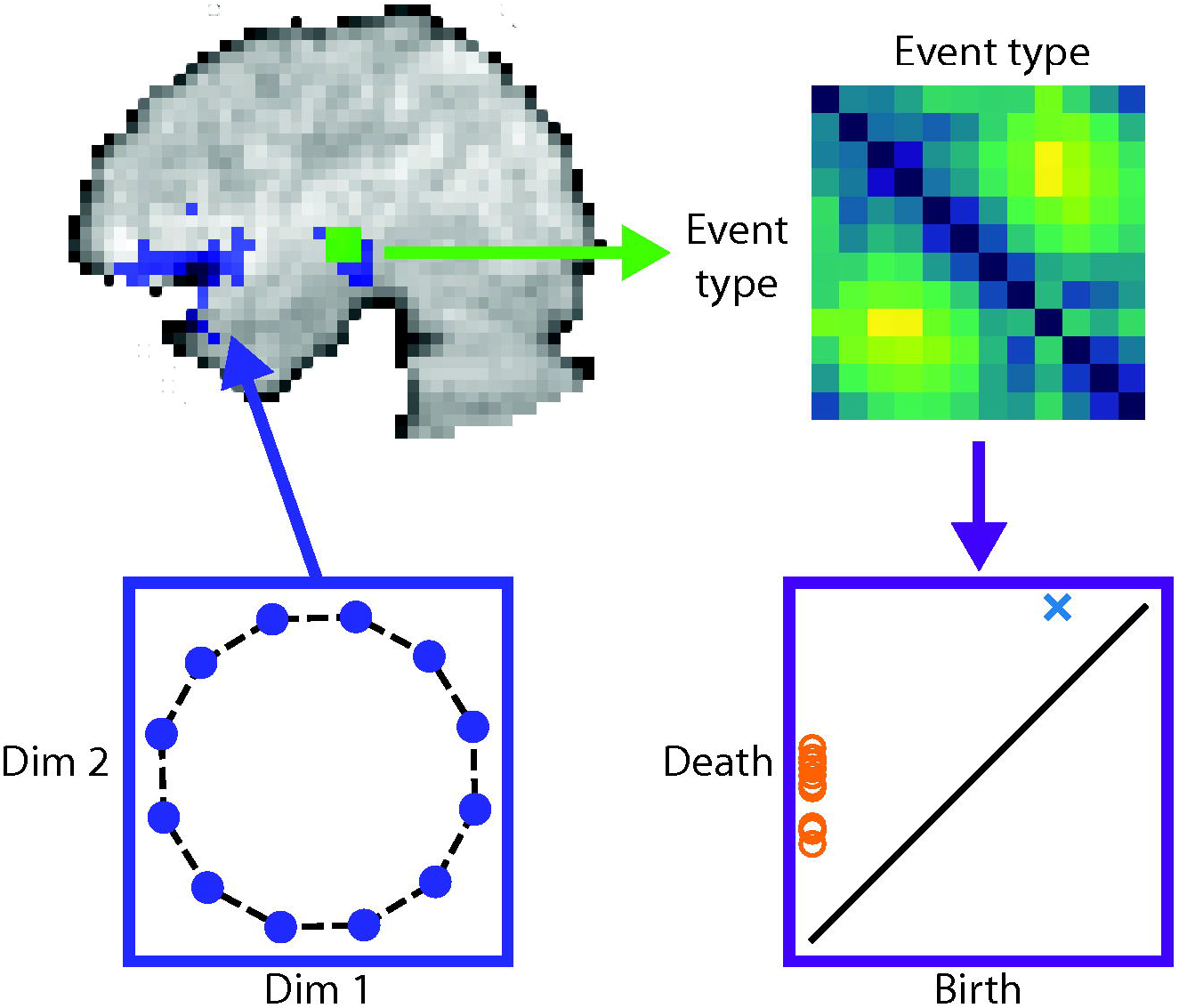
Application of persistent homology analysis to fMRI data. Lower left box shows a topological pattern of activity (N equidistant points lying on a circle in the Euclidean plane). The points in this set are embedded into an ROI in the brain, convolved with the hemodynamic response, and combined with noise (blue voxels in upper left image). A distance matrix is then calculated for each 3 x 3 x 3 searchlight of voxels by comparing the event-evoked activity for each event type (matrix at upper right). This distance matrix is the input to the persistent homology computation to create the barcode. The birth and deaths of cluster (orange circles) and loop (blue cross) features are plotted together (purple box at lower right). The persistence diagram is assigned to the central voxel that contributed to that searchlight.

To generate the temporal structure of the signal we took the N points sampled from the circle and treated them as event types that occur in an event-related design. An event type can be thought of as a unique stimulus type, like a landmark or face, that lies on the circle in high-dimensional voxel space. We generated designs in which different event types occurred in a trial sequence, split in to 5 runs. Each event type was repeated either 25 or 50 times per session, distributed equally throughout the 5 runs. This resulted in simulated session durations ranging from 60 to 180 minutes (more than 100 minutes would be a long fMRI experiment and might need to be completed across multiple sessions). For reference, 25 repetitions per event-type was equivalent to the usable fMRI data from Schapiro and colleagues (2013).

These events each lasted for 1s, and occurred 7, 9 or 11s apart with equal probability and in a randomized order. Although this delay is longer than what others like Schapiro and colleagues (2013) have used, this longer delay provides a cleaner measurement of the delayed, protracted nature of the brain’s response on which the fMRI signal relies (Burock, Buckner, Woldorff, Rosen, & Dale, 1998), and thus gives persistent homology a better chance to identify topological structure inherent in this signal.

Each event in this design evoked a characteristic pattern of activity across the voxels in the ROI. The magnitude of each voxel’s response to events was determined by the voxel’s embedding in the representation space. The maximum value in this embedding for each voxel (i.e. the coordinate furthest from the origin) was used to scale the response. Specifically the percent signal change of this coordinate was set to either 0, 0.25, 0.5, 0.75, or 1% and all other responses were then scaled proportionally relative to this maximum response. Percent signal change refers to the magnitude of evoked response relative to baseline fluctuations (Gläscher, 2009). In typical experiments, we expect to find between 0.25% and 1% signal change, depending on the cortical area and stimulus (Desmond & Glover, 2002; Rao, et al., 1996). The efficacy of simulating data with a given magnitude was validated in another study (Ellis et al., 2019).

Note, this simulation may be overestimating the power of these analyses because the participants being simulated have unrealistically consistent responses to the task. Although noise differences exist between simulated participants, the percent signal change that events evoke is all identical, which is not likely in most samples. Adding variability to the percent signal change in the simulation would add variability to the test statistics and diminish their significance. This was not implemented here since estimating the variability in percent signal change could not be done in systematically. Nonetheless, this is not expected to qualitatively alter the conclusions that can be drawn from this simulation.

Each voxel’s time course of evoked responses was convolved with a double gamma hemodynamic response function (Friston et al., 1998) to imitate the brain’s response to event onsets. These evoked responses were then added to the noise and Z-scored across time to complete the simulation of a run. For each combination of the conditions (number of event types, repetitions per run, and percent signal change), a new simulation was made for all runs and all participants.

### Analysis

To extract the evoked representation, a mass-univariate general linear model was performed on each run, following the standard approach with real fMRI data. To do this, all of the events for a given type were convolved with a double gamma hemodynamic response function (Friston et al., 1998) to predict the neural response, creating a coefficient for each event type at each voxel. These coefficients were then averaged across runs to create a voxelwise response to each event type. These averaged responses to each event type were then used in a BrainIAK (http://brainiak.org) searchlight analysis. A searchlight analysis involves iterating the same computation over a subset (“searchlight”) of voxels across the brain. The searchlight is a tensor of 3 x 3 x 3 x N voxels centered on every voxel in the brain, where N is the number of distinct event types. This searchlight size was chosen to suit convention. Figure S3 shows the results with a 7 x 7 x 7 x N searchlight. Compared to the results of the smaller searchlight, the results for this larger searchlight suggest that size may influence the false alarm rate.

Importantly, the signal we embedded was distributed arbitrarily across the 442 voxels in the ROI, but a searchlight can only contain a subset of these voxels. This was especially important since the shape of the ROI was not smooth or uniform. There were 2569 searchlights that contained at least one voxel in this ROI, and there were 325 searchlights out of the whole-brain that contained at least 10 voxels from the ROI. Nevertheless, since the signal being embedded into the high dimensional space was very low dimension (2D), this meant that the signal’s representation was highly redundant within this ROI. In other words, only a small sample of voxels was necessary to represent the signal structure. We suggest that this is biologically plausible: voxels in an ROI may represent different features or dimensions but any small sample of these voxels may be sufficient to differentiate representations, especially when the representations are distributed and thus likely to differ along many dimensions.

For each searchlight, we construct a 27 x N matrix, where each column of the matrix specifies the voxel pattern in the searchlight for one event type. Each column of voxels is Z-scored to standardize differences between the events. We then formed an N x N distance matrix by taking the Euclidean distance between each pair of columns of the 27 x N matrix. Events evoking similar patterns of activity within a given searchlight had a low distance between them, and vice versa.

A critical step in this analysis is that the columns of voxels are Z-scored; if this is not performed then the sensitivity to purely multivariate patterns is severely diminished (Davis & Poldrack, 2013; Walther, Nili, Ejaz, Alink, Kriegeskorte, & Diedrichsen, 2016). In this simulation, there was no mean difference in the evoked response; however, small, unsystematic differences in the average response to an event arise by chance. The size of these differences compared to those of the multivariate pattern that the stimulus evokes are sufficiently large to generate arbitrary differences in the Euclidean distance between events. Z-scoring the voxel activity within a condition eliminates these differences and thereby increases the sensitivity to differences in the multivariate pattern. Accordingly, we used the normalized Euclidean distance matrices in a persistent homology analysis of searchlights centered on every voxel in the brain. Note that, if mean differences between conditions are expected, for instance because one condition evokes a stronger response than the other (Davis & Poldrack, 2013), then Z-scoring may not be appropriate. An alternative approach is to use correlation as the distance metric, which also has the effect of normalizing mean differences.

We chose to use Euclidean distance here because our simulated signal ought to have no systematic mean differences between condition, and because it is a typical metric in the TDA community; however, in the neuroscience community correlation distance is often used (Kriegeskorte, Mur, & Bandettini, 2008). Figure S4 shows the results are qualitatively the same if correlation distance is used. These normalized Euclidean distance matrices were used as inputs into the persistent homology computation.

A full description of persistent homology is outside of the scope of the present work; interested readers are pointed to standard introductions to the topic (Carlsson, 2009; Ghrist, 2008). In brief, persistent homology takes as input a distance matrix and outputs an object called a barcode, which we explain below. The barcode is constructed from the data (i.e., the Euclidean distance matrix) by building a growing sequence of geometric objects called a *filtration.* We work with a standard filtration construction called the Vietoris-Rips complex (Carlsson, 2009).

The barcode is a list of intervals. We interpret the start of the interval (“the birth”) as the scale at which some hole in the filtration forms, and the end of the interval (“the death”) as the scale at which that hole closes up. The length of the interval (i.e., the difference between the birth and death) is called the persistence of that feature, and is usually interpreted as a measure of “importance” or robustness of the feature. Features that have small persistence are considered topological noise, whereas larger persistence is taken to reflect a meaningful topological feature (Cohen-Steiner, Edelsbrunner, & Harer, 2007). In fact, in topology we distinguish between holes of different dimensionality: a 0-dimensional hole corresponds to a cluster, a 1-dimensional hole is a loop, a 2-dimensional hole is a hollow sphere, etc. Each interval in the barcode is labeled by the dimension of the corresponding feature. One way to plot the output of persistent homology is as a persistence diagram, where the x-coordinates are births and the y-coordinates are deaths of features. We use different symbols to distinguish the dimensions of the features. Figure 1 shows a persistence diagram with a number of cluster features and a single loop feature.

Our analysis tests for the presence of the single loop feature that was embedded into the data as part of the simulation. We consider two test statistics of this signal: i) the number of loops identified in the barcode (which should be exactly 1); and ii) the persistence of the longest-lived loop (if any were identified).

To do so, we computed a barcode for every searchlight in the brain. This was done by first constructing the N x N Euclidean distance matrix specified above. This distance matrix was input in to BrainIAK-extra’s (https://github.com/brainiak/brainiak-extras) Python wrapper of the PHAT algorithm (Bauer, Kerber, Reininghaus, & Wagner, 2017) to perform the persistent homology computation. The barcode from this computation was assigned to the voxel in the center of the searchlight. This computation was performed on each searchlight in the brain. For computational efficiency, we looked only for cluster and loop features (i.e., persistent homology in dimensions 0 and 1). If persistent homology is an adequate tool for identifying the loop structure embedded in realistically noisy fMRI data, then within the signal ROI the persistent homology plots should have exactly one persistent (long-lived) feature; this should not be so for an analogous ROI that does not contain signal. For this control ROI, we used voxels on the symmetrically opposite side of the brain as the signal ROI, and did not have signal added to the simulation (also 442 voxels). These ROIs then serve as a basis of comparison to determine where there should and should not be high test statistic values (either the maximum persistence or the proportion of single loops).

To evaluate the reliability of the test statistics, we perform a whole-brain, unbiased statistical test that corrects for multiple comparison issues. Due to the nonnormal properties of the persistent homology metrics used here (i.e., maximum persistence is a value from 0 to infinity, or whether the barcode contains a single loop feature is a binary value), we subtracted the whole brain mean of each test statistic volume and then performed FSL’s ‘randomise’ function to compute the non-parametric reliability of test statistics across the sample participants (Winkler, Ridgway, Webster, Smith, & Nichols, 2014). Voxels surviving TFCE correction at p<0.05 are plotted for each metric and for each condition. Finally, we compared the proportion of voxels in the signal ROI that are significant to the number in the control ROI.

## Results

To visualize the analyses performed here, Figure 2 shows the data for the 12 event-types, 25 repetitions condition with different signal change parameters. Figure 2A shows the distance matrix from a single searchlight in a single subject in the signal ROI. This searchlight was used as an input into Figure 2B to produce multi-dimensional scaling plots (Shepard, 1962). Figure 2C shows the persistence diagram computed for this searchlight. These plots indicate that as the signal change increases, the fidelity with which the loop structure can be recovered is greatly increased.

**Figure 2.**
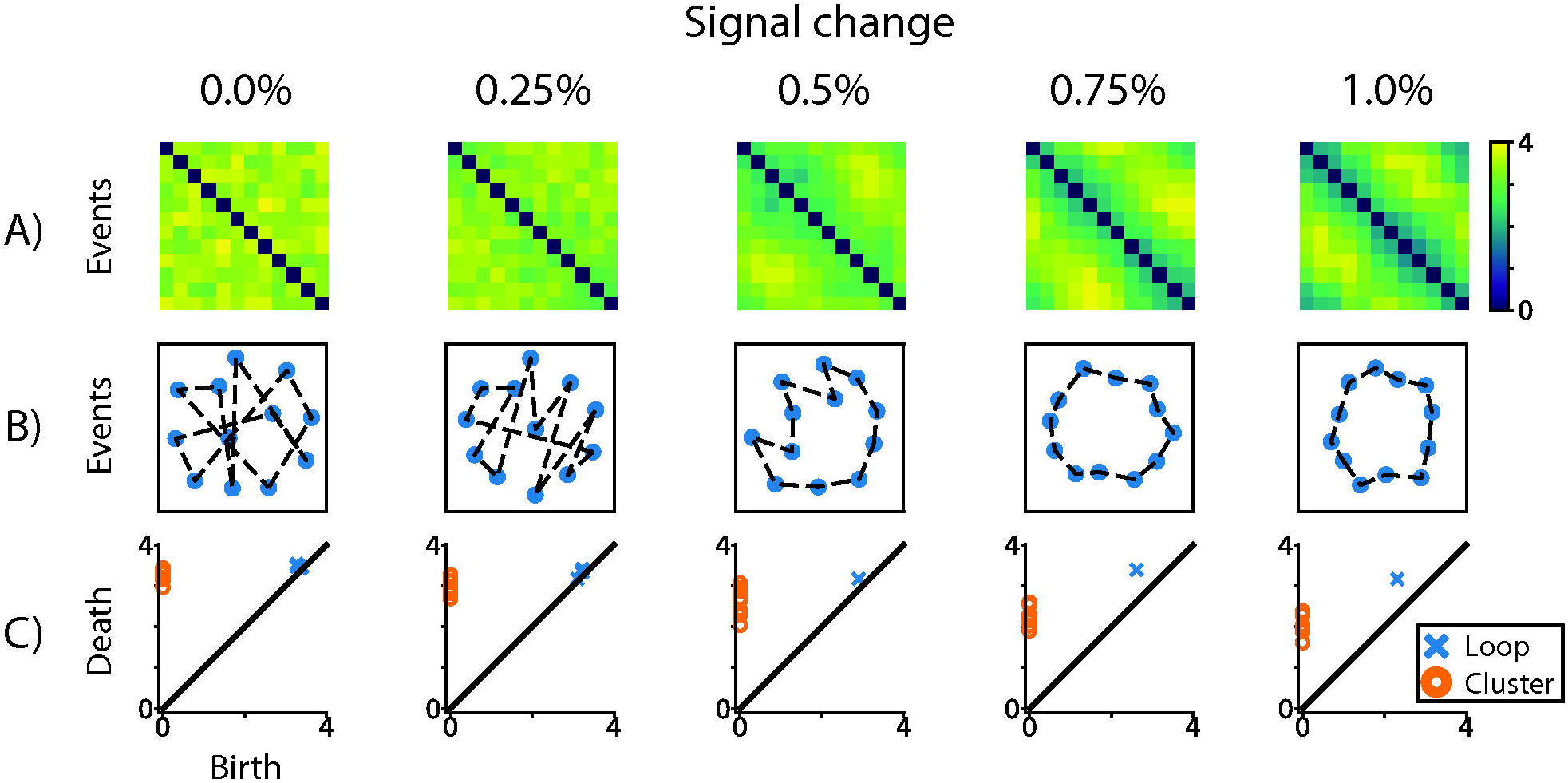
Data from example searchlight for different levels of percent signal change for the condition with 12 event types and 25 repetitions. (A) shows the event-by-event Euclidean distance matrices after voxels have been normalized within condition in this searchlight. (B) shows the MDS plots for these distance matrices. Lines connect events that should be adjacent in representation space. (C) depicts the persistence diagrams for cluster and loop features, derived from the distance matrices in (A).

To systematically quantify the evidence of a loop, we analyzed the size of the most persistent (or long-lived) loop feature. Figure 3A shows the mean persistence of voxels in the signal ROI subtracted from the mean persistence of voxels in the control ROI (i.e., voxels on the exact opposite side of the brain that did not contain signal). Each line represents a different set of experimental conditions (25 or 50 repetitions per session; 12, 15, or 18 event types), and each increment represents a different signal magnitude (0, 0.25, 0.5, 0.75, or 1% signal change). Figure 3A shows that with low (0.25%) signal that persistence initially dips in the signal ROI relative to the control ROI. With more signal, especially when there are more repetitions, the signal ROI shows greater persistence than the control ROI. Figure 3B shows the proportion of voxels in the signal ROI that are significantly reliable across participants. This shows a similar pattern as Figure 3A: the number of significant voxels only exceeds the baseline when there is at least 0.5% signal change, especially when there are more repetitions. Figure 3C shows a slice of the brain test statistic map for the 12 events, 25 repetition condition for each of the levels of signal change. Even at moderate signal there is evidence that the loop structure is expressed in the signal ROI. These findings suggest that it may be possible to recover a single, simple, clear topological signal with persistent homology in designs with a moderate evoked response and a large amount of data.

**Figure 3.**
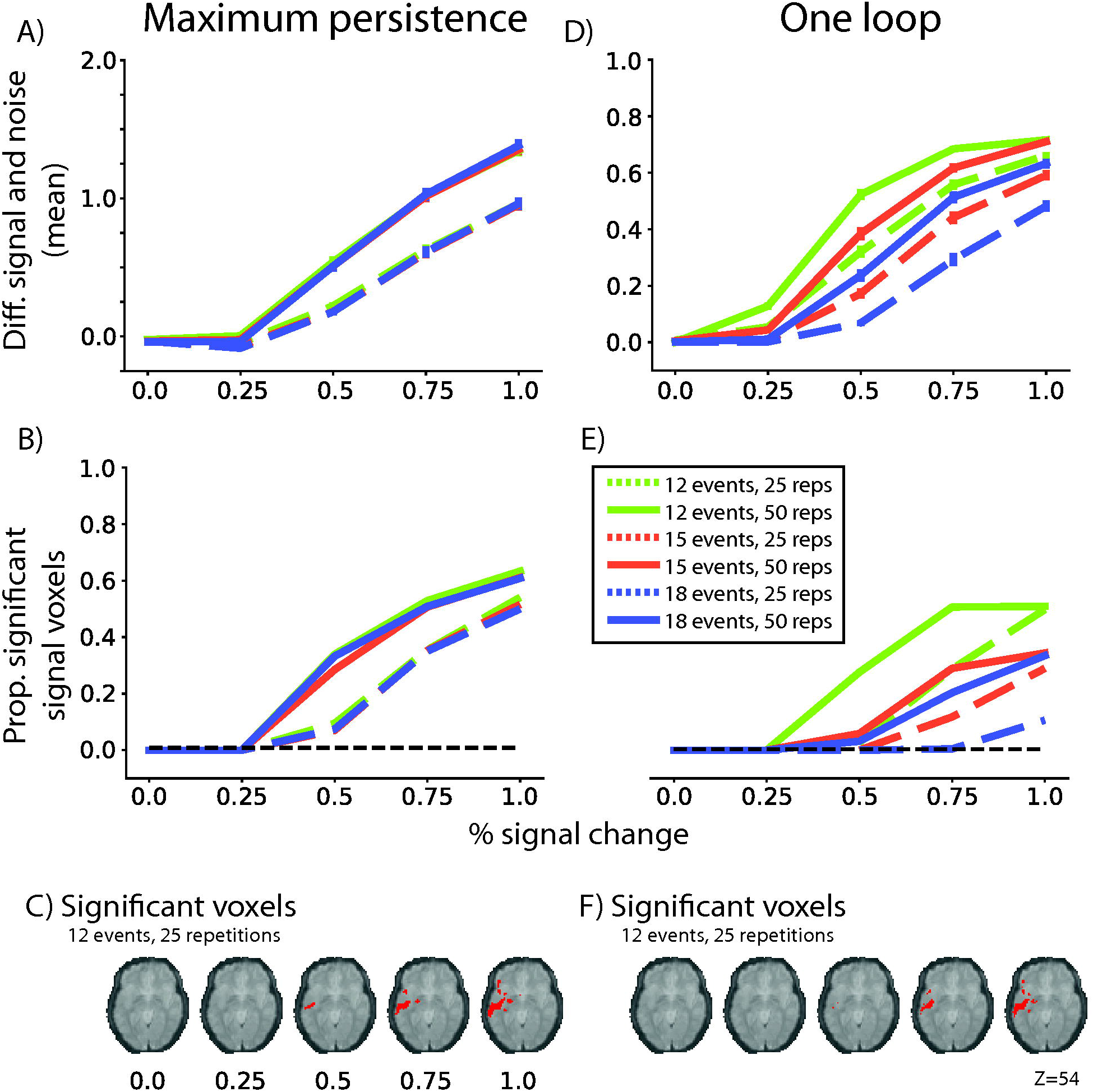
Results of persistent homology analysis of simulated event-related fMRI data. The left column shows the results using maximum loop persistence (i.e., length of the longest bar in the 1-D barcode) as the metric. (A) shows the difference in persistence in the signal ROI compared to the control ROI. Different colors depict numbers of event types (12, 15 or 18) and the different lines depict numbers of trial repetitions (25 or 50). (B) shows the proportion of voxels in the signal ROI that were significant. The black dashed line is the proportion of significant voxels in the control ROI. (C) shows in red the maps of voxels for which the maximum persistence is significantly greater than baseline at p<0.05 with TFCE correction for one condition of the experiment. Each brain represents a different percent signal change. The right column shows the equivalent analyses as (A-C) for the metric testing the proportion of voxels in the signal ROI exhibiting one loop feature (i.e. the 1-dimensional barcode with exactly one point). (D) shows the mean difference between signal and control ROIs. (E) shows the proportion of significant voxels in the signal ROI. (F) shows the significant voxels for each level of signal change. Error bars for (A) and (D) are standard error of the mean between participants.

Figures 2D-F shows the corresponding analyses for the test statistic evaluating whether there is only one loop feature (i.e. the associated 1-dimensional persistence diagram has exactly one point). Figure 3D shows that in some conditions there is evidence that the signal ROI can contain a loop structure even at the lowest signal change (0.25%). In the conditions with more event types or fewer repetitions, greater signal change was needed, though still in a reasonable range. Although these mean differences indicate evidence of a loop structure at low signal change, Figure 3E indicates that more signal is needed for the loop structure to be significantly reliable — only at 0.5% signal change or higher are single loops detected reliably. Figure 3F is similar to Figure 3C, and shows that even with moderate signal there is evidence that the loop structure is expressed in the signal ROI but nowhere else in the brain.

Additional analyses, reported in Figure S5, tested whether it was possible to improve upon the detection of single loops by taking into account “topological noise”. Topological noise refers to low persistence features that may emerge briefly in the presence of much larger and more robust high persistence features. We can ignore these features by setting a threshold on the length of persistence that is sufficient to warrant inclusion relative to the length of the next longest feature. By introducing this threshold, the rates of finding 1 loop increase for both the signal and control ROI. Hence it is necessary to find an optimal threshold for which the control ROI is kept low and the rate in the signal ROI is high. There are indications that when using a low signal (0.25%) it is possible to find greater loop evidence in the signal ROI relative to the control ROI when using a ratio threshold between 1.4 and 2.6. However, the optimal threshold differs depending on the number of event types and thus is difficult to set a *priori.* Hence, topological thresholding may hold value but determining a principled way of applying it to neural data requires further investigation.

## Discussion

Here we report simulation analyses evaluating the usefulness of persistent homology in recovering topological structure in event-related fMRI data. Specifically, we used fmrisim to simulate a realistic pattern of noise in fMRI data, and inserted a specific topological feature (a single loop) into some voxels. We calculated the persistent homology of all voxel-centered searchlights in the brain and evaluated the extent to which the embedded loop was reliably identified by persistent homology. We showed that persistent homology can extract structure from a sample of participants collected with a realistic event-related fMRI design, especially when there is moderate signal and few event types, with many repetitions of each type.

We used two different metrics to evaluate the topological structure inserted into the data: (1) maximum persistence of a loop feature and (2) whether there is only one loop feature present in the barcode. We found generally consistent results between these methods; however, there were some circumstances in which the evidence conflicted. When the signal strength was low and there were few repetitions of each event, the measure of maximum persistence was actually lower in the region of the brain containing signal compared to a control region. At the same time, the proportion of searchlights that contain a single loop in the signal ROI was greater than the control ROI, even though this was not significantly reliable across participants. There are other circumstances where the best metric is less clear. If a larger searchlight is used, our supplemental analyses suggest that the proportion metric may be more susceptible to noise than maximum persistence. It is hoped that this simulation protocol could help to investigate these nuances and other novel designs.

The pattern of results showed that topological signal was best recovered when (i) the signal strength was high, (ii) there were numerous repetitions per event, and (iii) there were fewer events in the circle. The benefits of high signal strength and increased repetitions were expected; however, it was not anticipated that fewer event types would make it easier to extract the representation of the loop. This phenomenon may emerge from the greater risk that a single noisy event will disrupt the topology of the entire representation. That said, with too few events, the topological structure cannot be formed.

Although we were successfully able to use persistent homology to recover signal, it is important to consider a number of limitations that were exposed in this simulation. Firstly, the distance metric used, as well as the preprocessing done on this metric, is extremely important: it is possible to entirely miss signal in the brain by doing the wrong type of preprocessing on the data. We strongly advise using metrics that normalize voxel activity within condition when the differences between conditions are expected to result from multivariate patterns. Secondly, the amount of data needed in these analyses (60 to 180 minutes) is substantial and may preclude certain types of experiments. Thirdly, the topological structure that we tested is simple and described by relatively few points. Hence for more complicated topological features, such as community structure (Schapiro et al., 2013), “figures of 8”, interlocking rings and branching structures, it is likely that persistence analysis will have even lower power. Nonetheless we believe the pipeline we set up provides an opportunity to test the viability of extracting different topological structures from event-related fMRI.

It is also possible that other experimental designs may profit from the use of TDA to identify mental representations. For example, designs using more naturalistic stimuli (e.g., movies or narratives; Hasson, Nir, Levy, Fuhrmann, & Malach, 2005; Yeshurun et al., 2017) generate large quantities of data (in which each time point can be treated as a node on a graph) with strong evoked responses. Such data may be conducive to analysis using TDA. Moreover, persistent homology is only one type of TDA; it may be that other types of TDA, such as Mapper (Singh, Mémoli, & Carlsson, 2007), can also recover topological structure in fMRI data.

In sum, we have shown that topological structure embedded in realistic simulations of fMRI data can be identified and we have characterized conditions under which persistent homology is highly effective. We hope that the insights and tools introduced here can guide researchers in discovering the topological structure of neural representations.

## Acknowledgements

Thank you to N. Malek for comments on an earlier draft. We thank the editors and reviewers for their helpful feedback. We especially thank one of the anonymous reviewers who went above and beyond what is expected from reviewers and substantively improved the manuscript.

## Contributions

C.T.E., M. L., B. K. & J.D.C. designed the project. C.T.E., M. L., G.H.P., B. K. developed the analysis. C.T.E. performed the analyses. C.T.E. wrote the initial draft of the manuscript. All authors reviewed the manuscript.

