## Supplementary Material for "Feasibility of Topological Data Analysis for event-related fMRI"

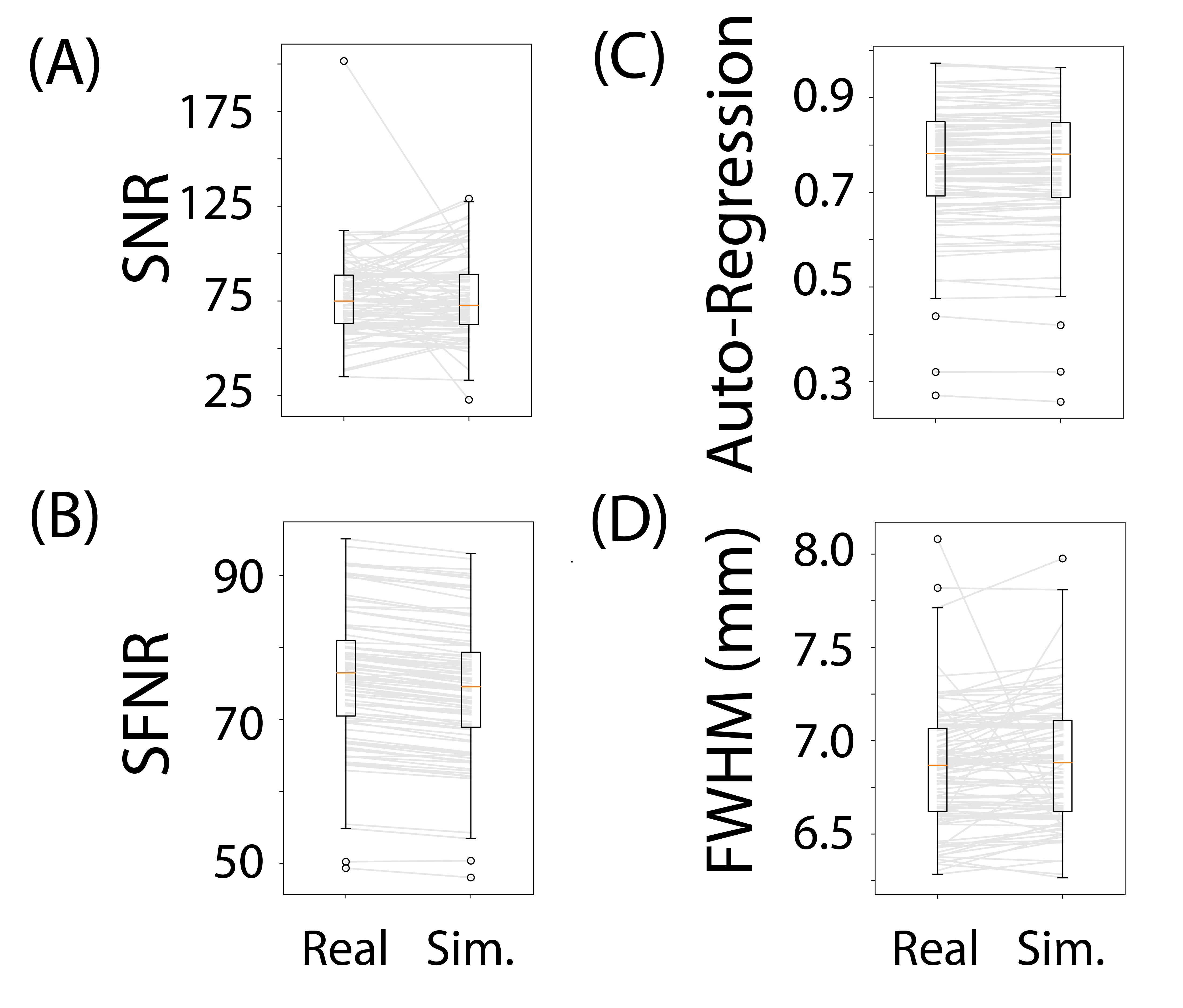

*Figure S1*. Boxplots of noise parameters from real data and fitted simulated data. For each participant and run, 10 simulations of their real data were generated and their noise parameters estimated. These noise parameters were then averaged within participant/run and plotted, and compared with the noise parameters from their corresponding real data. The grey lines connect the participant/run parameter value, the ellipses refer to outliers. The noise parameters tested were (A) Signal-to-Noise Ratio (SNR), (B) Signal-to-Fluctuation-Noise Ratio (SFNR), (C) Auto-Regression, (D) Full-Width Half-Max (FWHM).

*Interpretation:*

The critical feature of these plots is that for the most part the simulated parameter values are within the range of the real values.

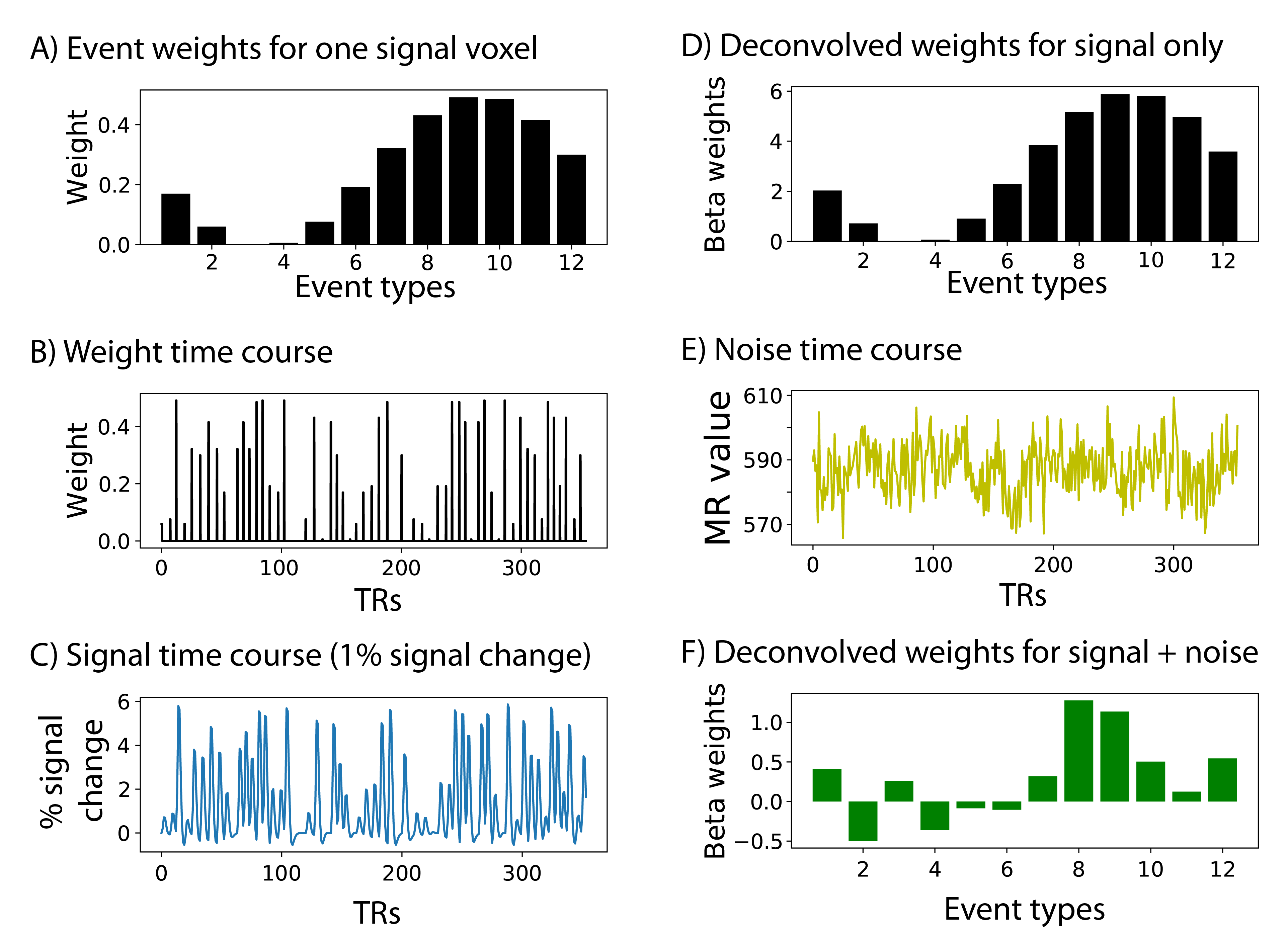

*Figure S2.* The signal transformation involved in simulation and preprocessing. A) shows the weights for a specific voxel for each event type (after orthonormal transformation and rescaling all values to be positive). High numbers mean this voxel will respond more strongly when this event occurs. B) shows the hypothesized neural response over the course of a run. Each response reflects the onset of an event, and the height is determined by the weight for that event. C) is the expected response to the events shown in (B). This is the result of a convolution with the double-gamma hemodynamic response. The voxel’s maximum response across the run is rescaled to 1% signal change. D) shows the result of a univariate analysis of this voxel without the addition of noise. Specifically, this shows the beta weight for each event type by comparing the voxel’s response to a hypothesized response for each event type separately. Importantly, because there is no noise, this graph has the same shape as (A) but is scaled to 1% signal change. E) is the time course of the noise for this voxel, before any signal is added. Note that the mean of this time course is approximately 600, meaning that a 1% signal change corresponds to approximately 6 units of increase. F) This is the same analysis as (D) except that it is of a voxel after both signal and noise have been added, and then the data was Z-scored in time. Evidently, there is substantial noise relative to what signal was inserted; nevertheless the pattern is largely consistent with (A). These beta weights for each voxel are then averaged across runs and used in subsequent analyses to compute the similarity between events.

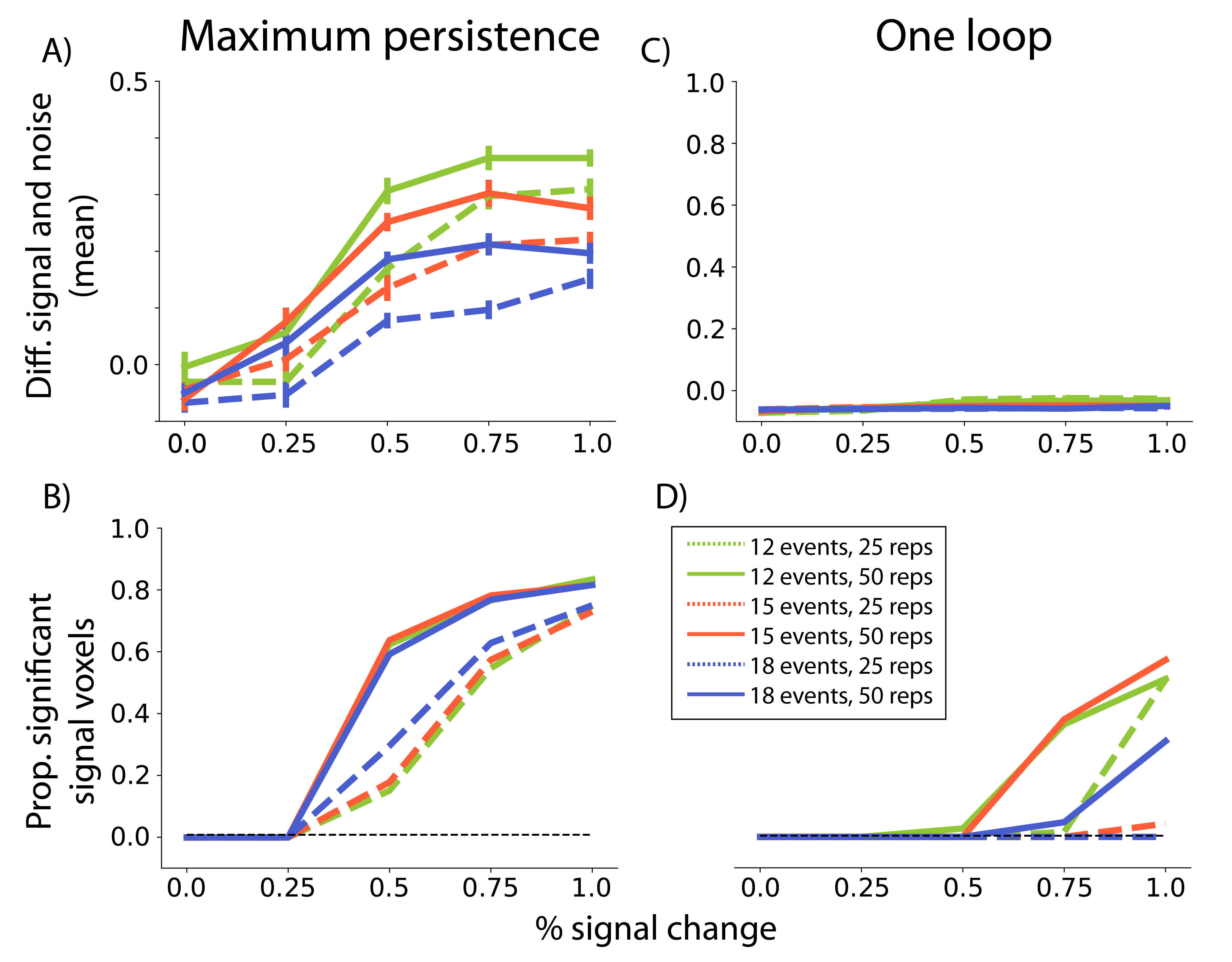

*Figure S3.* Equivalent analyses to Figure 3 when using a searchlight size of 7 x 7 x 7 x N. Refer to the caption of Figure 3 for more details. Note: the underlying simulated data is the same here as reported in the main text, the only difference is which searchlight radius was used.

*Interpretation*

Increasing the searchlight size improves the maximum persistence metric but diminishes the single loop metric. Particularly striking is that in (C) the signal ROI is below the control ROI but (D) shows that some voxels are still significant. Based on our follow-up analyses we observed two potential reasons that account for this: (1) the false alarm rate increases with larger searchlights because the chance of noise data to form a persistence diagram with many loops diminishes. For instance, in the 0.0% signal change, 25 repetition and 12 event condition, the number of loops per searchlight, averaged across the whole brain of a single participant, is 10.0 for a 3 x 3 x 3 searchlight and 5.6 for a 7 x 7 x 7 searchlight. (2) multiple loops often form in searchlights along the edges of the signal ROI, and since the searchlight is large, this happened more often. The reason that some voxels are still significant is that those voxels that are not on the edge are reliable across participants, thus making them significant; whereas the searchlights that form 1 loop by chance are not reliable across participants.

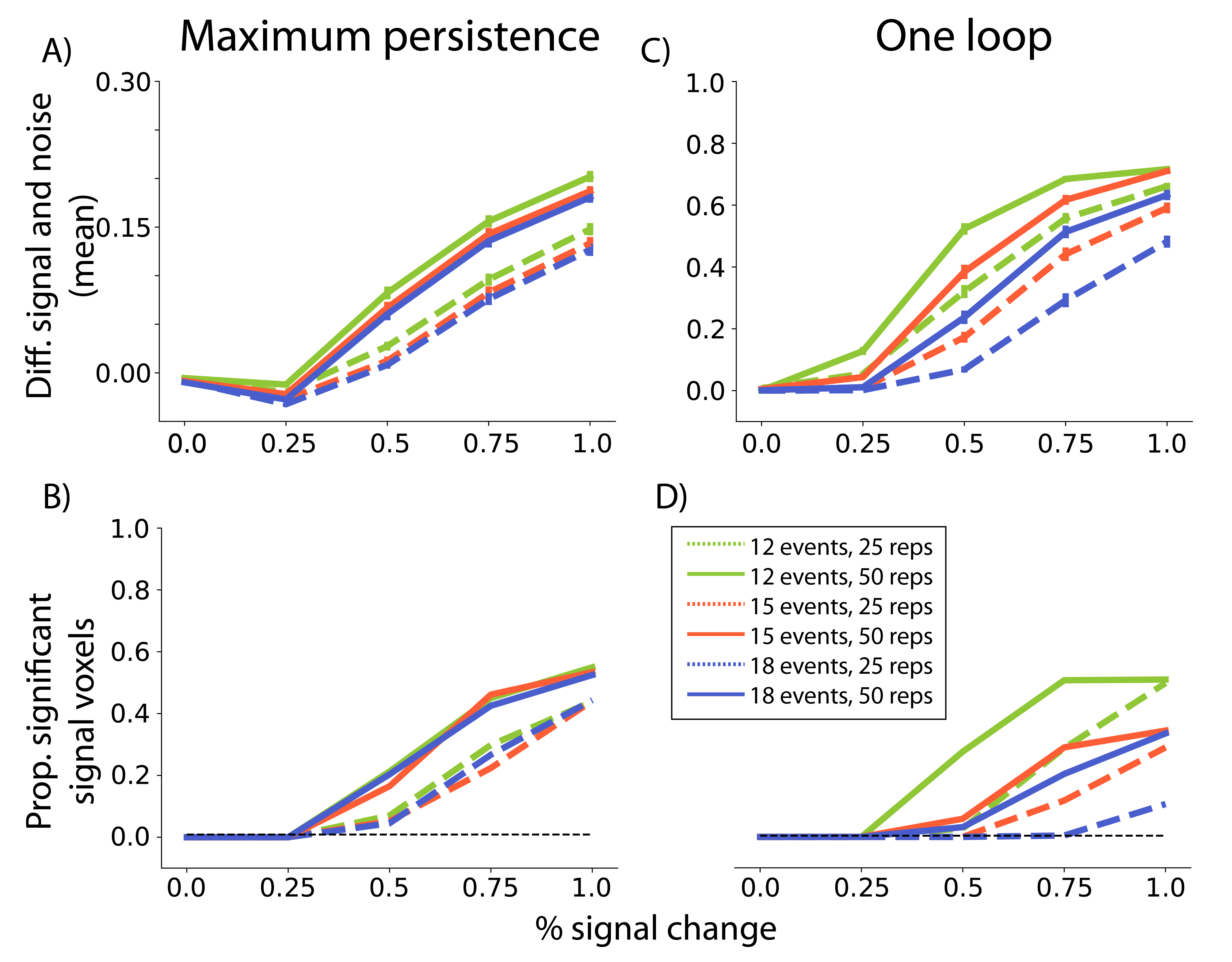

*Figure S4.* Equivalent analyses to Figure 3 when using correlation as the distance metric. Refer to the caption of Figure 3 for more details. Note: the underlying simulated data is the same here as reported in the main text, the only difference is which distance metric was used.

*Interpretation*: There is very little difference between using the normalized Euclidean metric reported in the main text and the correlation metric depicted here. This is because the Euclidean distance matrices and correlation matrices that are used as the basis for persistent homology are very similar. The scale is different and this contributes to some small deviations of the maximum persistence metric from Euclidean distance; however, this scale change has almost no effect on the proportion of voxels with a single loop.

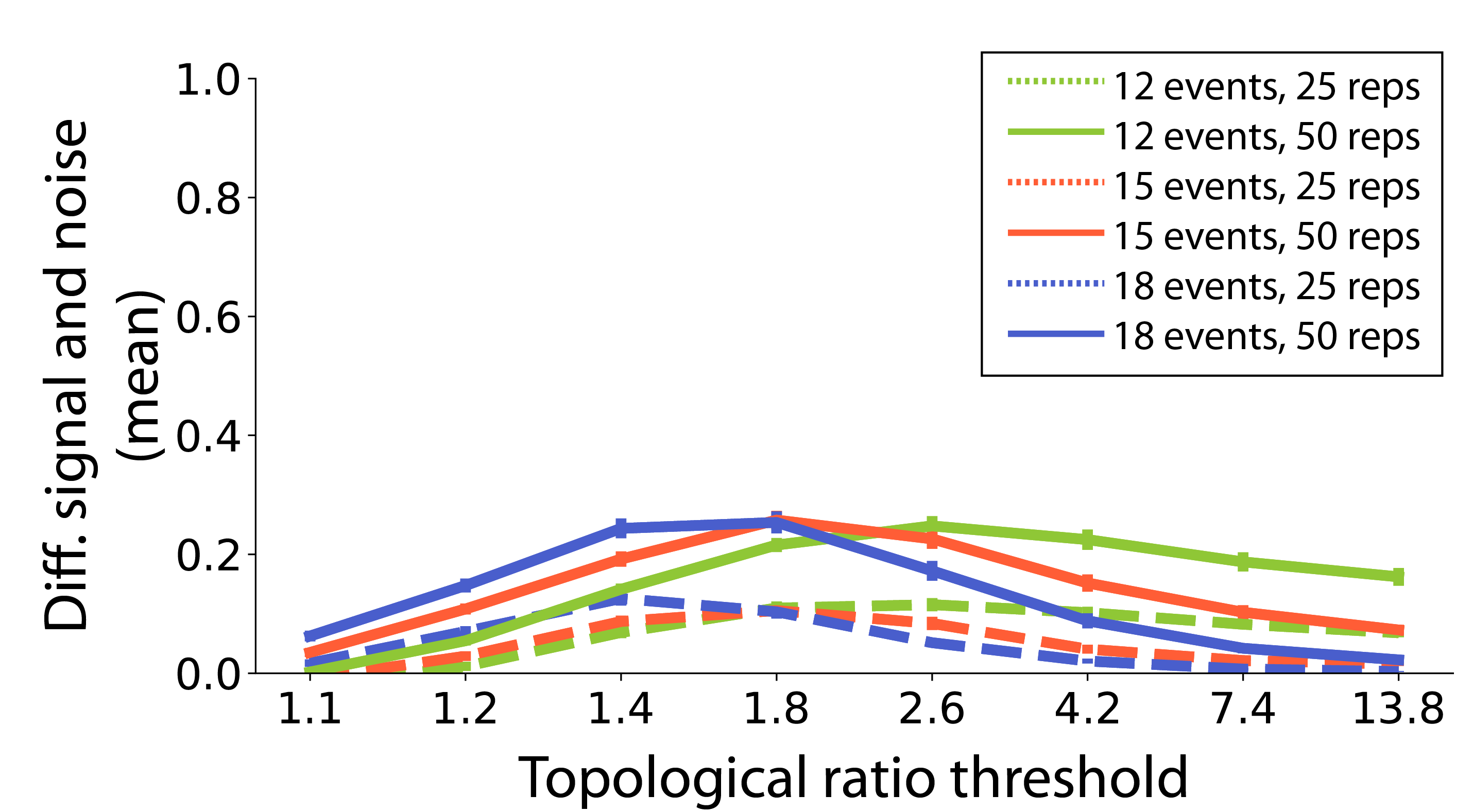

*Figure S5.* Difference in proportion of voxels with 1 loop in the signal ROI compared to the control ROI (similar to Figure 3D). This plot shows the low signal (0.25%) data for each condition, searching over different topological ratio thresholds. The ratio threshold refers to the ratio of lifespans between the longest- and second longest-lived loops that is necessary to count a persistence diagram as having 1 above threshold loop (e.g., a ratio of 2 means the longest-lived loop must be twice as persistent as the next longest-lived loop to be counted). If there is only one loop, this is automatically a “hit”, and if there are zero loops, this is automatically a “miss”. Note: the underlying simulated data is the same here as reported in the main text, the only difference is what topological threshold was applied to the data.

*Interpretation:* Although Figure 3D indicates that at 0.25% signal change there is little difference between the signal and control ROI for 15 and 18 events, when a topological threshold is set appropriately there is some evidence of signal. In particular when the threshold is 1.4-1.8 the difference between the signal and the control ROI is maximal for these conditions. Critically, a different threshold, around 2.6, is appropriate for the conditions with 12 events.
